# Left-right asymmetry of the microminipig brain

**DOI:** 10.64898/2026.01.25.700707

**Authors:** Yuga Fujiwara, Kaichi Yoshizaki, Rio Mikoshiba, Takahito Arai, Na Wang, Azusa Seki, Masaki Takasu, Naokazu Goda, Satomi Chiken, Atsushi Nambu, Yoshiaki Shinohara

## Abstract

Left-right asymmetry of the brain is well recognized in various animals including C. elegans, drosophila and zebrafish. In primates, most of the brain studies describe side of the brain. However, in spite of huge amounts of accumulating rodent studies on neuroscience, most of rodent studies do not distinguish the brain side. The pig brain is considered to occupy an intermediate position between primates and rodents in terms of structural complexity and brain function. Moreover, the numbers of studies using genetic manipulation of pigs are drastically increasing. So, we investigated microminipig (MMP) brain mesoscopic anatomy focusing on left-right differences of its morphology. Here, we show the anterior cingulate cortex, perirhinal cortex, and cerebellum of male and female MMPs, are structurally asymmetrical. The cerebellar vermis, paravermis is tilted from the midline and the consequently the cerebellar cortex exhibits asymmetrical morphology. The anterior cingulate gurus exhibited protrusion and invagination toward the midline on the right and left side, respectively. The left perirhinal lobe exhibited distinct patterns of cortical gyration between left and right side. These data demonstrate that MMPs are one of the suitable model animals for investigating cerebral and cerebellar asymmetry.

## Introduction

Mounting evidence suggests that brain laterality is commonly observed in many kinds of animals including mammals ^1^, zebra fish^2^, drosophila^3^, snails and nematodes ^4,5^. Among them, functional brain laterality is well known in primates such as humans^6,7^ and monkeys^8^. For example, hand motor control and language are dominant in the left neocortical hemisphere^9^. More circuitry-based mechanisms of brain asymmetry have been analyzed in the rodent hippocampus and auditory cortex. Left-right differences of the mice hippocampal synapses with differential receptor allocation have been reported ^10,11^, which are genetically determined ^12^. In addition, postnatal stimuli influence on hippocampal functional laterality and gene expressions in rats ^13^. Asymmetries of the neuronal circuitry are also reported in the mouse auditory cortex ^14,15^. Developmental speed of the auditory cortex is reported to be influenced by animal sex and laterality of the mouse brain ^16^. Though human functional neocortical asymmetry has gained interest of general audience, molecular basis of neocortical asymmetry is largely unaddressed.

One of the emerging model animals for life science study is a pig. Being familiar animals as livestock, pigs are also known to be highly intelligent and recognize themselves in their image in the mirror. Moreover, pigs can navigate with mirror images ^17^. Because of the brain size and anatomical complexity, pig brains are easier to translate into human brain research compared with rodents’ brains. The numbers of genetic manupilation studies in pigs are increasing in recent years, particularly for clinical applications such as interspecies organ transplantation^18–20^. Indeed, the use of pigs in biomedical research is gradually increasing, and more than 60,000 pigs were used annually in the European Union ^21,22^. Another notable advantage of pigs is a low psychological barrier for using them as experimental animals. As behavioral and psychological experiments using domestic pigs are technically difficult owing to their body size, microminipigs (MMP) have gained attention due to their small body size and experimental suitability. Thus, pigs provide valuable experimental models that bridge the physiological and anatomical differences between primates and rodents, enabling the extrapolations of research outcomes into human brains^23^.

In the present study, we investigated MMP brain mesoscopic morphology by 60 μm Nissl-stained coronal slice preparation combined with 7T MRI imaging. Left-right asymmetry of the brain was clearly visible in the cerebellum. Though asymmetric morphology of the cerebellum has been well recognized in cats ^24^, animal experiments using cats are increasingly difficult because of animal welfare. Since right cerebellar cortex is implicated in social recognition and language processing in humans ^25,26^, MMP cerebellar asymmetry is expected to be a novel animal model for studying cerebellar functional asymmetry.

The neocortical lobes, including the anterior cingulate gyrus and perirhinal lobe of MMP brain also exhibited asymmetrical morphology. The right anterior cingulate gyrus showed protrusion to the midline whereas left side is concaved. The asymmetrical structure of the anterior cingulate gyrus covers most of longitudinal (rostral – caudal) axis of this gyrus. In addition, an asymmetrical structure was observed in the perirhinal cortex, and the lobe exhibited distinct neocortical gyration patterns between left and right side. Using slice specimen of the neocortex, we verified asymmetry of above three areas and also analyzed the cytoarchitecture of the asymmetrical brain areas. Both anterior cingulate gyrus and perirhinal cortex has reciprocal projections between hippocampal formation^27–30^, and both areas are thought to work with hippocampus and play important roles in memory formation^31,32^. Therefore, elucidating the asymmetical circuitry of these two brain areas would give us cues for understanding hippocampal asymmetry.

Our findings indicate that swine brain also has apparent asymmetrical structures like primates, and we propose that with advances in genetic manipulation technologies in pigs, MMP are expected to be one of the valuable model animals for investigating brain asymmetry.

## Materials and Methods

### Microminipigs (MMPs)

Two male and one female MMPs were used for the present study. All animal experiments were conducted in accordance with the Guideline for the Proper Conduct of Animal experiments of the Science Council of Japan, and were approved by the Ethical Review Committee of the University of Yamanashi Graduate School of Medicine [Approval No. A5-18] and the Ethical Review Committee of Fuji Micra Inc (Shizuoka, Japan). Two male MMPs were purchased and one female MMP was kindly donated from Fuji Micra Inc.

### Rearing of MMPs

Rearing of MMP was performed as described elsewhere ^33^. MMPs were purchased from Fuji Micra Inc and kept in a controlled room with a temperature of 24 ± 3°C and a humidity of 50–70%. Lighting in the room was set on an 11 h light/13 h dark cycle starting at 08:00 h. The MMPs were fed with MMP pellets (3% BW, Marubeni Nisshin Feed Co, Ltd, Tokyo, Japan), a feed specially developed for MMPs, which comprise total digestible nutrients (TDN, >74%), crude protein (>13%), crude fat (>2.0%), crude fiber (>8.0%), crude ash (<1.0%), calcium (>1.1%), and phosphorus (>0.9%), and had free access to water.

### Preparing of the brain

The animals were sedated by intramuscular injection of xylazine (0.09ml/kg). Anesthesia was conducted by 3-4% isoflurane and then maintained with isoflurane at 1.5-2%. During surgical operation, isoflurane concentration was increased to 4%. Animals were perfused via the bilateral common carotid arteries with 0.1M phosphate buffer (PB), followed by a fixative solution containing 4% paraformaldehyde in 0.1M PB (pH 7.4). The brains were post-fixed and cryoprotected in a sucrose solution (∼30%).

### MRI acquisition and analysis

One female MMP was used for MRI analyses. The fixed brain was scanned using a 7-tesla whole-body MR scanner (Magnetom 7T, Siemens) with a 24-channel multi-array receive coil (Takashima Seisakusyo Co. Ltd. ^34^). The brain sample was placed in a container filled with PBS (0.01 M). Most of the air bubbles present within the sample were removed prior to scanning by applying negative vacuum pressure overnight. Twelve T2-weighted (T2w) images with 0.25 mm isotropic resolution were acquired using a sampling perfection with application-optimised contrast using different angle evolutions (SPACE) sequence (field of view (FOV): 80 x 80 mm; repetition time (TR): 6000 ms; echo time (TE): 451 ms; flip angle (FA): 120 deg), and sixteen T1-weighted (T1w) images with 0.4 mm isotropic resolution were acquired using a magnetization-prepared rapid gradient-echo (MPRAGE) sequence (FOV: 76 x 80 mm; TR: 2500 ms; TE: 2.41 ms; inversion time: 1100 ms; FA: 5 deg: average of 2 scans). The total scanning time was approximately 12 hours.

Each of the T1w and T2w images were averaged and rigidly aligned to a minipig MRI template ^35^ using FSL ^36^ with manual adjustments. The images were then resampled to 0.25-mm isotropic resolution, and bias-corrected by using the square root of the product of T1w and T2w images ^37^.

### Histological analyses

Two male MMPs were used for histological analyses. Coronal sections of the brain were prepared using an HM440E Sliding Microtome (ThermoFisher). For Nissl staining, each section was stained with 0.1% cresyl violet, sequentially exposed to 95% EtOH + 10% acetic acid for 5 min, 95% EtOH for 1 min, 100% EtOH for 1 min and subjected to xylene clearing. Images were then acquired using a microscope, BX63F (Evident) and BZX800 (KEYENCE).

## Results

### A reference point of the MMP skull for anterior-posterior coodinate for this study

First, we set reference points in the skull to make anterior-posterior coordinates of the MMPs. To perform electrode implantation and tracer injection into MMP brain efficiently in the future studies, easy-to-use standard points of the MMP skull for stereotaxic surgery are necessary. However, the bregma and the lambda, standard references for rodent brain surgery, is hard to visualize and sometimes indiscernible in adult MMPs (orange and purple circles in Fig.1a, respectively). In addition, the bregma is located near the anterior extreme of the MMP brain, the point is inadequate for reference for the whole MMP brain. To circumvent these problems, we newly set a standard point (point-FS (point of full width of the <u>s</u>kull)), which is the crossing of the line where width (transverse length) of the skull is the maximum and midline of the brain (red circle in Fig. 1a). In the lateral view, the bilaterally extended line from the point-FS was located along the dorsal margin of the temporal fossa and posterior to the orbit (Fig.1b). In the posterior view, the occipital bone, including the external occipital protuberance, the mastoid processes of the temporal bone, the posterior ends of the zygomatic arches, and the region surrounding the foramen magnum were observed (Fig.1c). Subsequently, histological analyses were conductd with this point-FS as a reference.

**Figure 1.**
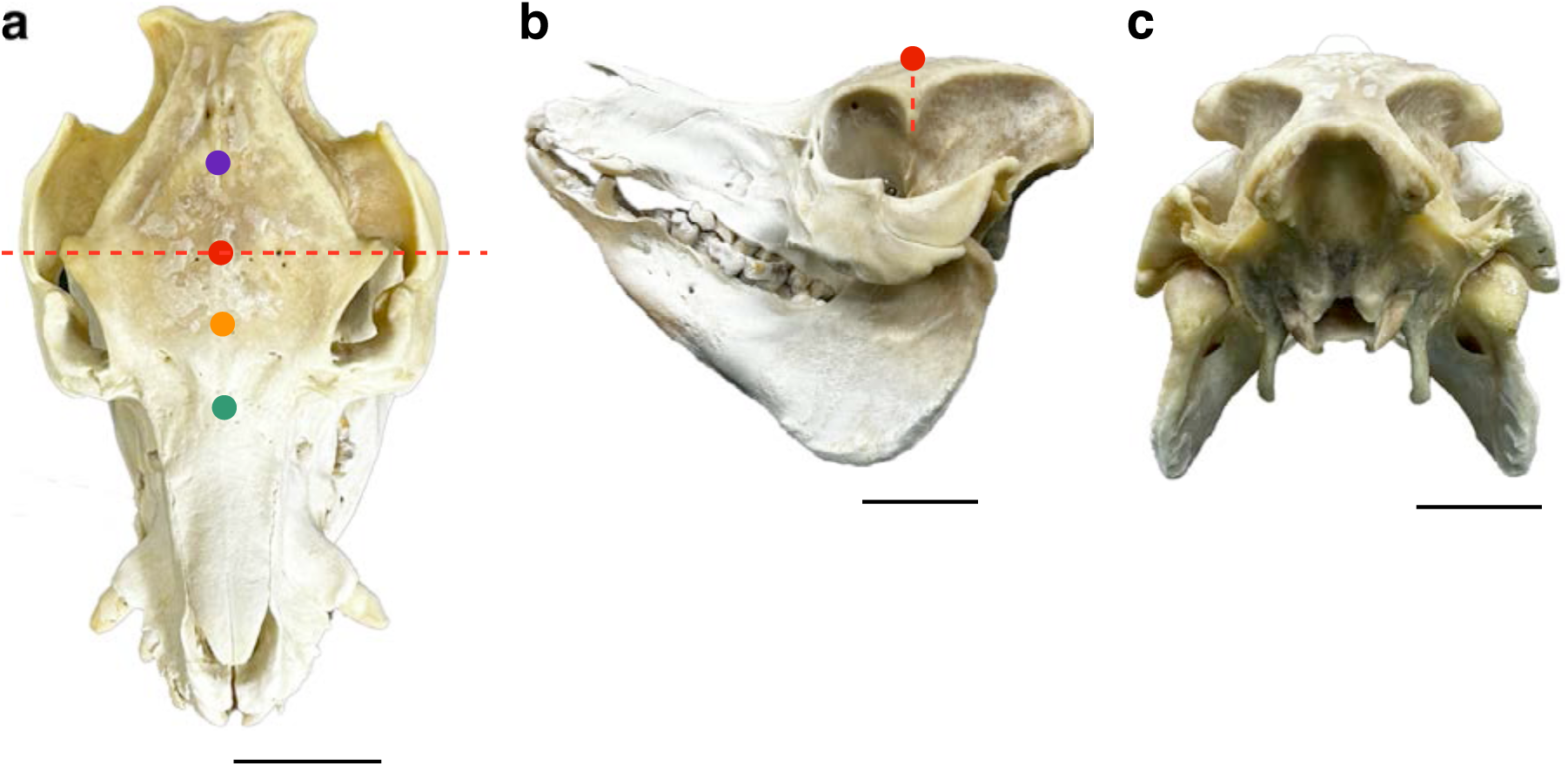
MMP skull overview. (a) A dorsal view of the MMP skull. Anatomical landmarks, such as bregma (orange circle) and lambda (purple circle) are hardly visible in adult MMP skull. Point-FS (red circle) is defined as the intersection point of widest line of parietal bone (dashed red line) and the midline. Green circle, nasion. (b) A lateral view of the MMP skull. (c) A posterior view of the MMP skull. Scale bar: 5 cm.

### External appearance of the MMP brain

Next, we removed MMP brain from the skull cavity and investigated the exterior appearance of the brain. A lateral view (left-side view) (Fig. 2a), dorsal view (Fig. 2b), and ventral view (Fig. 2c) of the brain are exhibited, respectively. The brain of MMP contains many gyri and sulci, we defined the major neocortical lobes according to Göttingen minipig ^38^ (Fig.2e). The lateral view (Fig. 2a) exbibhits large cerebrum (Cer), smaller cerebellum (Cbel) and lower part of the brain including the olfactory bulb (OB), olfactory tubercle (Tu), and brainstem (BS). A prominent Sylvian sulcus was visible, which connects to presylvian sulcus anteriorly and rhinal fissure posteriorly, separating the cerebrum from the brainstem.

**Figure 2.**
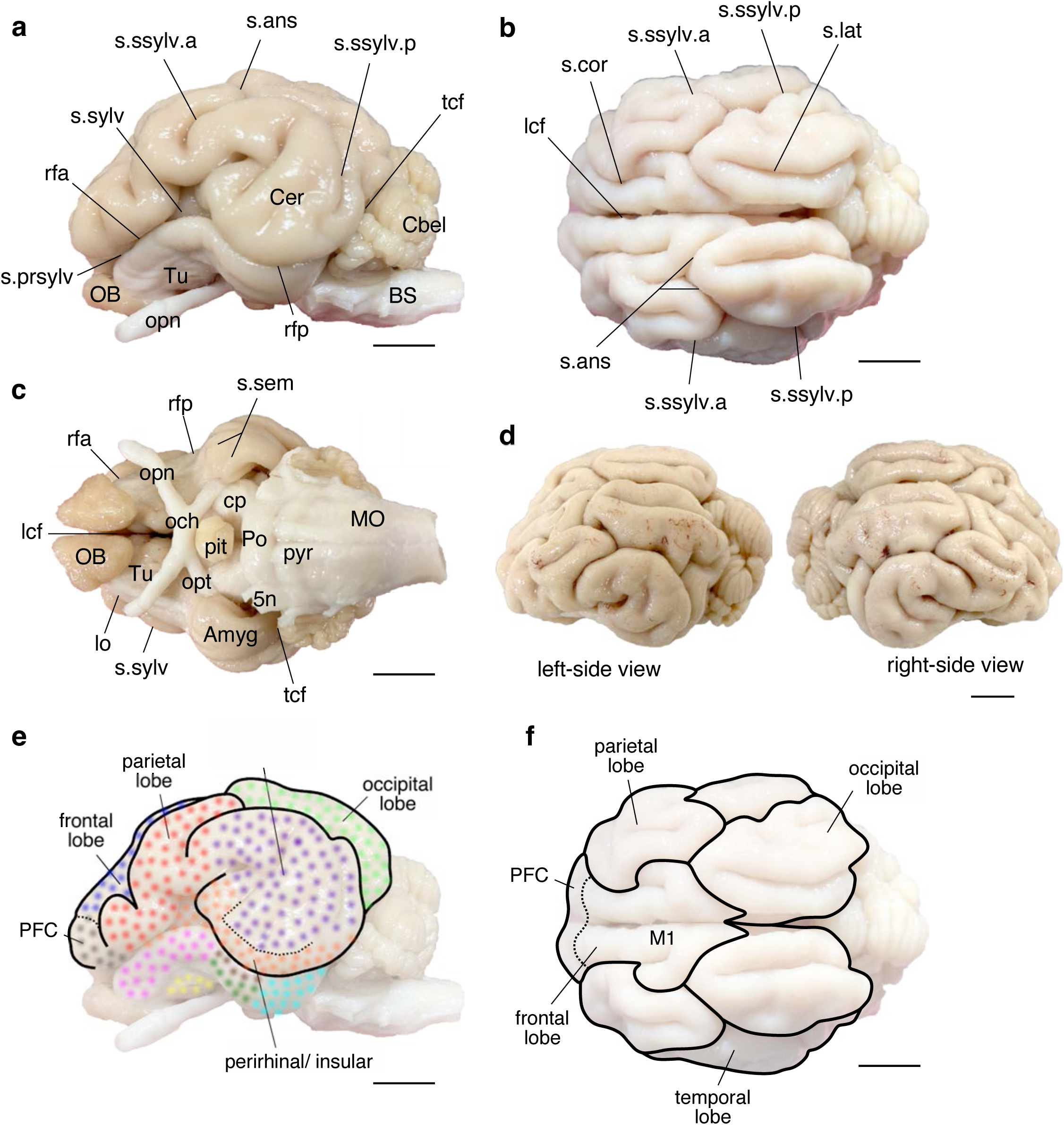
Macrophotos of the MMP brain. (a) Lateral view, the anterior (rfa) and the posterior (rfp) parts of the rhinal fissure separates the cerebrum into a dorsal gyrencephalic part and a ventral subrhinal part. (b) Dorsal view, the ansate sulcus (s.ans) form an important landmark. (c) Ventral view, the semiannular sulcus (s.sem) marks the position of the amygdala (Amyg). (d) Lateral view from right and left side. (e) Photograph of panel A (lateral view) with auxiliary lines overlaid for reference, depicting the segregation of the MMP telencephalon into lobes and some of their major subareas. Note the location of the perirhinal/ insular cortices in the anterior and ventral regions of the temporal lobe. (f) Photograph of panel b (dorsal view) with auxiliary lines overlaid for reference, depicting the segregation of the MMP telencephalon into lobes and some of their major subareas. Note that the visible subareas of the frontal lobe, *e.g.* the prefrontal cortex (PFC) and the motor cortex (M1), are indicated separately. *Amyg*: amygdala, *BS*: brain stem, *Cbel*: cerebellum, *Cer*: cerebrum, *cp*; cerebral peduncle, *lcf*: longitudinal cerebral fissure, *lo*: lateral olfactory tract, *MO*: medulla oblongata, *OB*: olfactory bulb, *och*: optic chiasm, *opn*: optic nerve, *opt*: optic tract, *pit*: pituitary gland, *Po*: pons, *pyr*: pyramis, *rfa*: rhinal fissure anterior part, *rfp*: rhinal fissure posterior part, *s.ans*: ansate sulcus, *s.cor*: coronal sulcus, *s.lat*: lateral sulcus, *s.prsylv*: presylvian sulcus, *s.sem*: semiannular sulcus, *s.ssylv.a*: suprasylvian sulcus anterior part, *s.ssylv.p*: suprasylvian sulcus posterior part, *s.sylv*: sylvian sulcus, *tcf*: transverse cerebral fissure, *Tu*: olfactory tubercle, *5n*: trigeminal nerve. Left and right sides are indicated as L and R, respectively. Scale bar: 1 cm.

The dorsal view of the brain (Fig. 2b) shows Cer and Cbel. The lateral cerebral fissure (lcf) lies in the midline of the Cer and divides the left and right neocortical hemispheres. The frontal cortex is observed in the anterior and central part of the brain, which is separated from parietal and occipital lobe by the coronal sulcus (s.cor) and the ansate sulcus (s. ans), respectively (Fig. 2b and 2f). The motor cortex (M1) occupies a large portion of the frontal cortex. The prefrontal cortex (PFC) was located at its rostral extremity. The parietal lobes were located in the lateral part to the frontal lobe, and the lobes can be identified as bilateral protrusions of the lateral brain. The ventral border between the parietal lobe and the lower brain part such as Tu was the anterior part of the rhinal fissure (rfa). The occipital lobes are located in the center of the posterior Cer with a large groove of the lateral sulcus (s.lat). The lobe is separated from temporal lobe by the posterior part of the suprasylvian sulcus (s.ssylv.p). Temporal lobes are located in the lateral portion of the middle-posterior brain, lateral to the occipital lobe and posterior part of the parietal lobe. The border between the temporal lobe and the lower brain part such as the BS and the amygdala (Amyg) is the posterior part of the rhinal fissure (rfp).

The ventral view (Fig.2c) indicates OB, Tu, Amyg, and brain cerebral peduncle (cp), pons (Po), and medulla oblongata (MO) and cranial nerves. The Cbel is separated by transverse cerebral fissure (tcf). We found that the dorsal view clearly indicates that the caudal vermis of the cerebellum is skewed to left-side (Fig. 2b). We confirmed this asymmetrical structure of the cerebellar vermis by the left-side view and right-side view of the brain of another MMP (Fig. 2d).

### MRI imaging of fixed MMP brain

MRI images of a MMP brain were obtained using a 7-tesla scanner (Fig. 3). Examples of coronal (T1w: Fig 3a; T2w: Fig. 3b), sagittal (T1w: Fig 3c; T2w: Fig. 3d), and horizontal (T1w: Fig 3e; T2w: Fig. 3f) images are shown, respectively. The complex neocortical structures of the MMP were clear and we identified large lateral ventricles around hippocampus and in the center of bilateral striatum.

**Figure 3.**
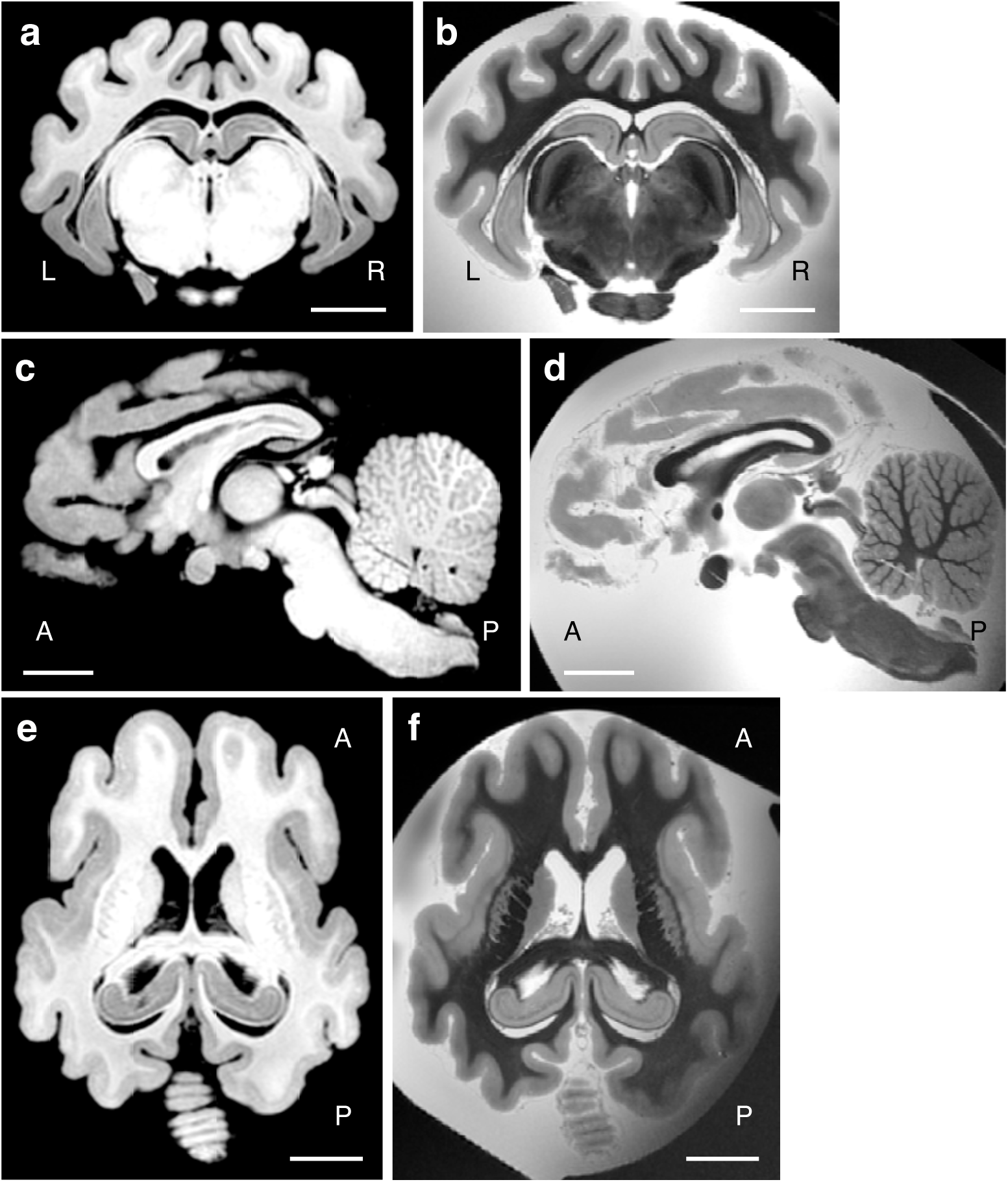
Representative MRI images of the MMP brain. Representative coronal (a, b), mid-sagittal (c, d) and horizontal (e, f) sections of T1w (a, c, e) and T2w (b, d, f) MRI images. Left and right sides are indicated as L and R, respectively. Anterior and posterior side are indicated as A and P, respectively. Scale bar: 1 cm.

### MRI images of the coronal section exhibit asymmetry in the cerebrum and cerebellum

To examine the MRI images in greater detail, we selected five coronal slices. The positions of the images are indicated as dashed lines in the lateral and dorsal views, respectively (Figs. 4a and 4b). Figs. 4c and 4h show T1w and T2w MRI images of the most anterior portion of the neocortex, respectively, where each hemisphere was separated into three gyri, the frontal and the medial and lateral part of the parietal lobe. In addition, each hemisphere is clearly divided into left and right hemispheres. At the ventral part of the slice, the olfactory bulbs are visible. Within the olfactory bulbs, lateral ventricle olfactory part can be identified as hollow circular shape structures.

**Figure 4.**
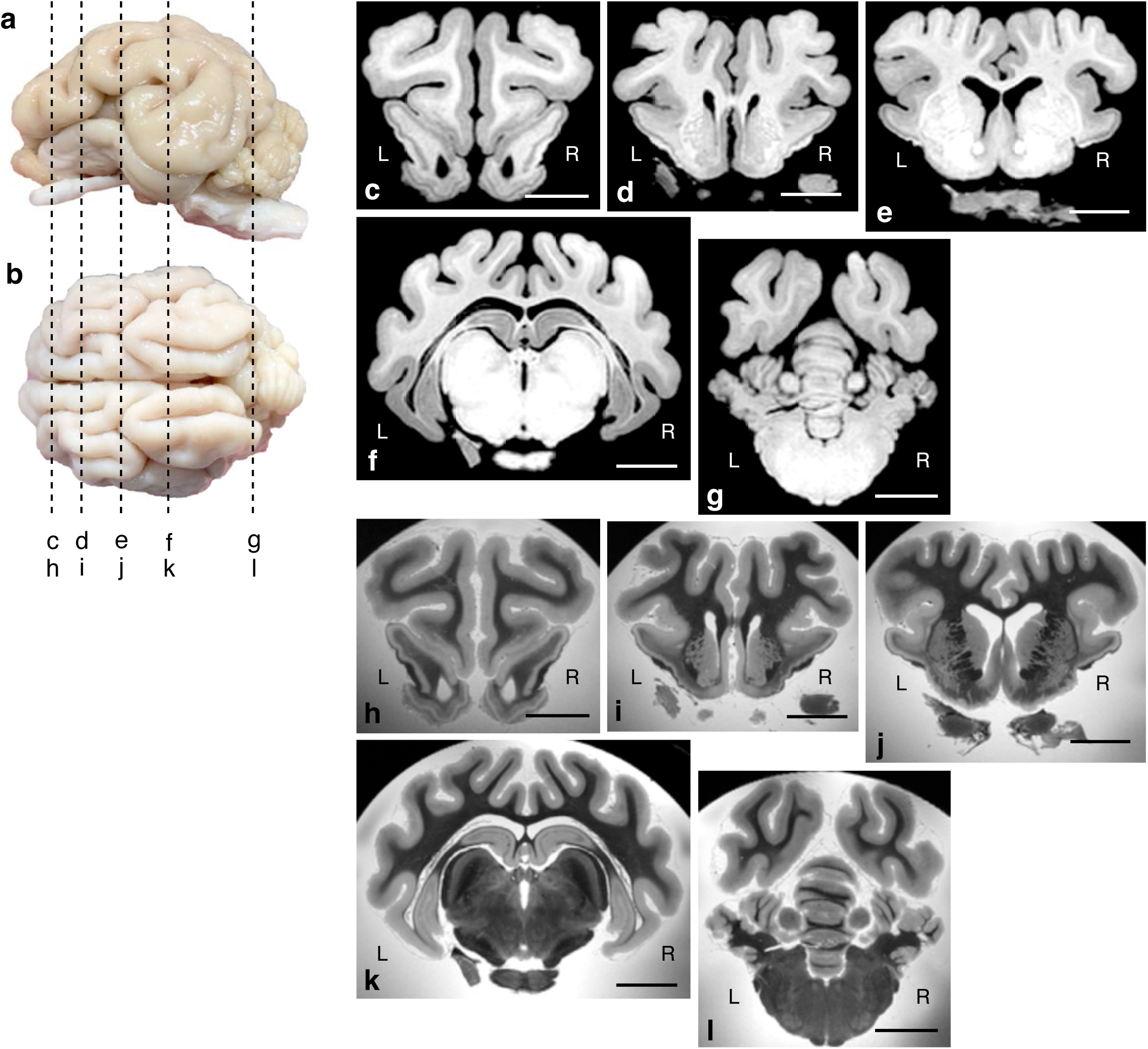
MRI images of the coronal section. Lateral (a) and dorsal (b) views of MMP brain. Vertical dashed lines indicate the approximate plane of coronal levels used for subsequent MRI analyses along anterior-posterior axis. (c-l) Representative coronal sections of T1w (c-g) and T2w (h-l) MRI images corresponding to vertical dashed lines. Left and right sides are indicated as L and R, respectively. Scale bar: 1 cm.

MRI images in Figs. 4D (T1w image) and 4I (T2w image) indicates more posterior position of the brain. The anterior parts of the striatum are clearly visible and lies ventral to the neocortex. The left and right neocortices are connected with the corpus callosum which are located at the center of the lateral ventricles. The frontal and parietal lobes were visible in the dorsal part of the image, Notably, we found a protrusion toward the midline was observed in the right neocortex, corresponding to the cingulate gyrus.

Figs. 4e and 4j show more posterior section, and the protrusion of the right hemisphere becomes more prominent, whereas the left side exhibits an invagination of the cingulate gyrus just above the corpus callosum.

At the position of Figs. 4f and Fig. 4k, the occipital cortex occupied the central dorsal region of the neocortex, laterally franked the temporal lobes. At this level, asymmetry of the neocortical gyri becomes less apparent. In the ventral part of the neocortex, lateral ventricles, and both dorsal and ventral part of the hippocampus are clearly visible. In the ventral central region of the section, the BS structures such as thalamus and subthalamus, occupy the space.

In the most posterior part of the brain (Figs. 4g and 4l), the Cbel and MO are identified. As suggested in Fig. 2d, the central part of the MRI image, which corresponds to the vermis, is asymmetrical.

### Asymmetrical structure of the MMP cerebellum

As indicated by Fig. 2 and Fig. 4, the asymmetrical structures of the vermis of the cerebellum is apparently visible. So, we sought to examine more detailed structures of the asymmetrical morphology of the cerebellum. By setting the point-FS as a reference, we visualized MMP brains at a histological slice level. In addition, we set the horizontal level by the line between the line of ventral extrememes of OB and BS (i.e. bottom of the brain), and obtained coronal slices by using this horizontal line. We examined the coronal slice morphology at +51.48 mm from point-FS in Fig. 5a.

**Figure 5.**
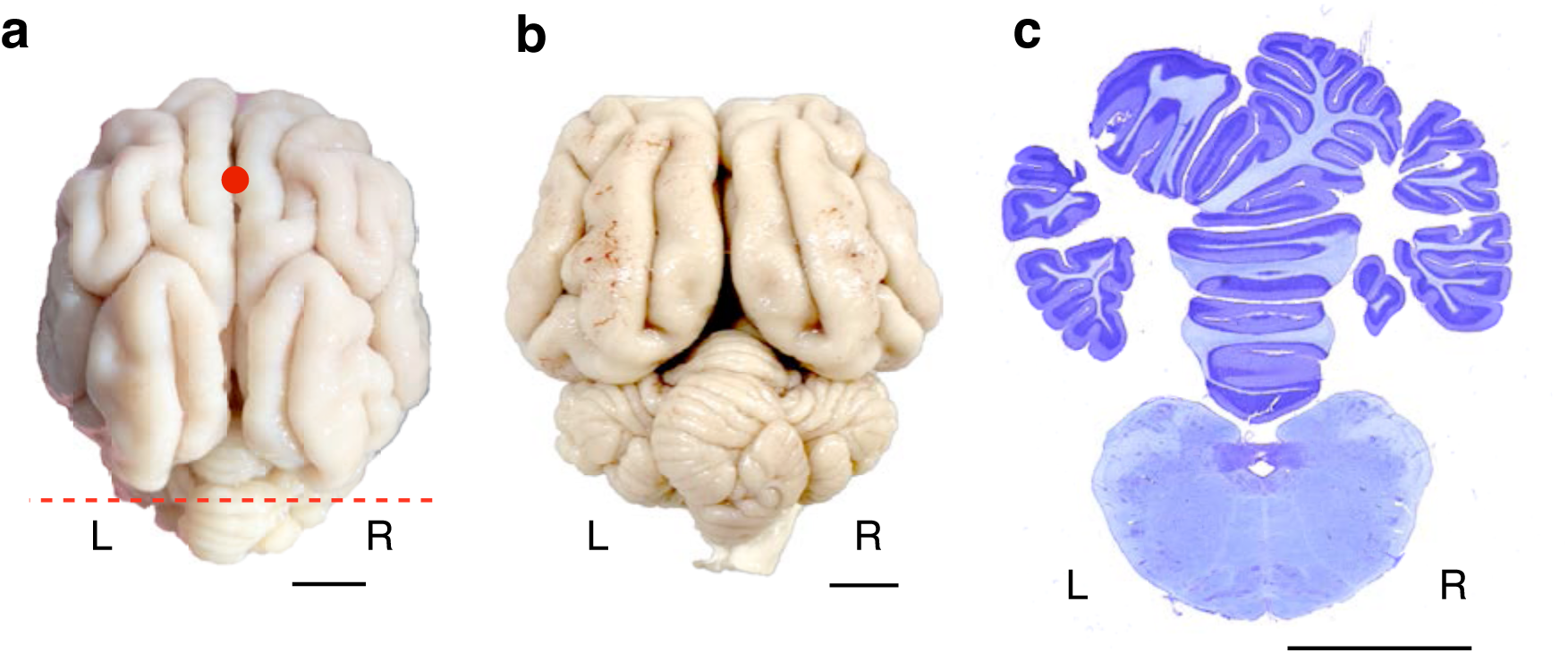
Asymmetry in the white matter of the cerebellum. (a) Dorsal view of the MMP brain with reference point, point-FS (red circle). The red dotted line (+51.48 mm posterior from point-FS) indicates the position of the coronal section corresponding to panel C. (b) Oblique posterior view of the MMP brain. (c) Coronal brain section at +51.48 mm posterior from point-FS in panel A. Left and right sides are indicated as L and R, respectively. Left and right sides are indicated as L and R, respectively. Scale bar for external appearance: 1 cm, Nissl staining of whole brain: 1 cm.

The coronal section of the cerebellum in the dotted line was shown in Fig. 5b. The lateral lobes of the cerebellar cortex are symmetrically located to left and right side of the brain. However, the section of the vermis of the cerebellum is Y-shaped, and the while matter bundles in the midline are running to right and dorsal direction. Interestingly, more ventral side of the vermis, this asymmetrical structure is tapered, and the medulla oblongata is symmetrical to body axis.

Next, we examined cerebellar cortical structures by using MRI sagittal sections (Figs. 6a, 6b and 6c). Figs. 6b and 6c indicates left side and right side sagittal sections of Fig. 6a, respectively. As shown in Fig. 6b (left, ML: −4.5 mm) and Fig. 6c (right, ML: +4.5 mm), cerebellar structures exhibited distinct structures between left (Figs. 6b and 6d) and right (Figs. 6c and 6e) sides.

**Figure 6.**
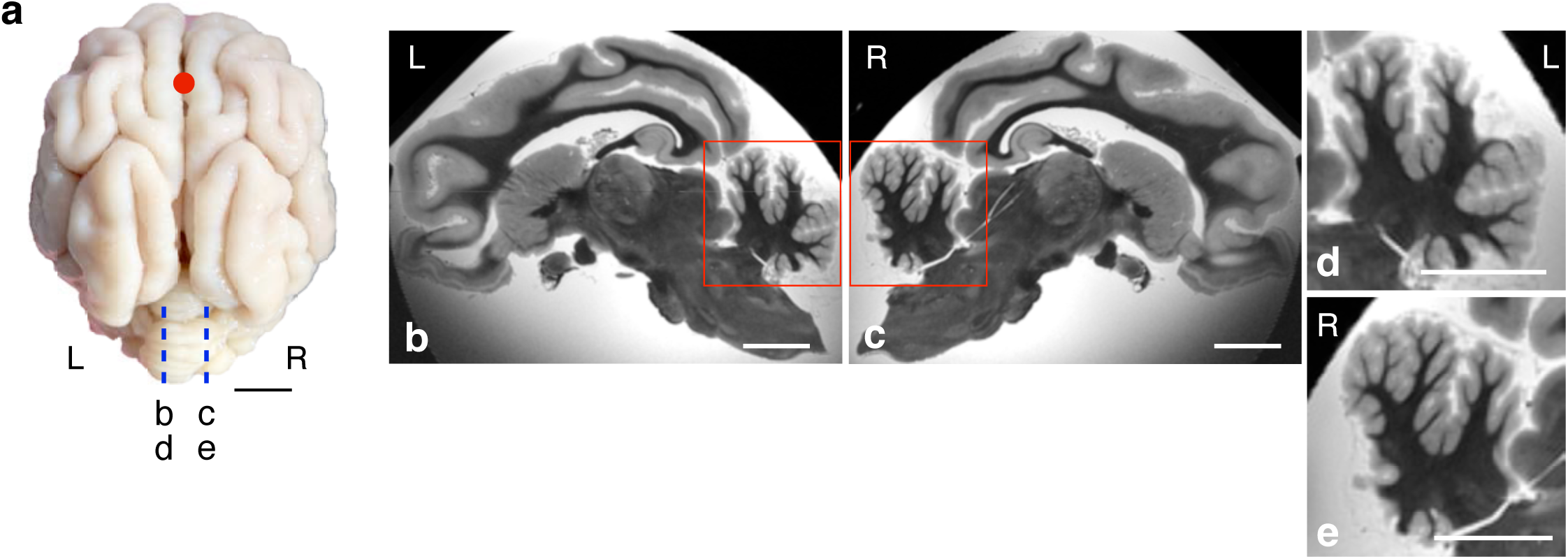
Asymmetrical structures of the cerebellum. (a) Dorsal view of the MMP brain with reference point, point-FS (red circle). Left and right of the sagittal sections are indidated by dotted lines (left: b and d; right: c and e, respectively). Representative sagittal section of T2w MRI images taken 4.5 mm lateral to the midline on the left (b, d) and right (c, e) hemispheres, respectively. (d, e) High magnification of the cerebellar cortex from the left (d) and right (e) hemispheres. Left and right sides are indicated as L and R, respectively. Scale bar: 1 cm.

### Nissl staining images at the frontal lobe

Given the asymmetrical morphology of the cerebellum, we next sought to investigate cerebrum structures. We examined the coronal section at −5.04 mm from point-FS (Fig. 7a). As indicated in Fig. 7b, the neocortex and the olfactory bulbs are separated. In addition, two hemispheres were separated by midline, resulting in four brain subsections. The neocortex of the images comprises the two major gyri, the frontal lobe and parietal lobe as viewed from the medial aspect, and the olfactory bulb contains lateral ventricles in the central part. The Nissl staining of the frontal lobe revealed an agranular cortex characterized by large layer V pyramidal cells (Fig. 7c), indicating that this region corresponding to the motor cortex. In the parietal lobe, a granular cortex with prominent layer V pyramidal cells was observed (Fig. 7d), suggesting that this region corresponding to the parietal association cortex.

**Figure 7.**
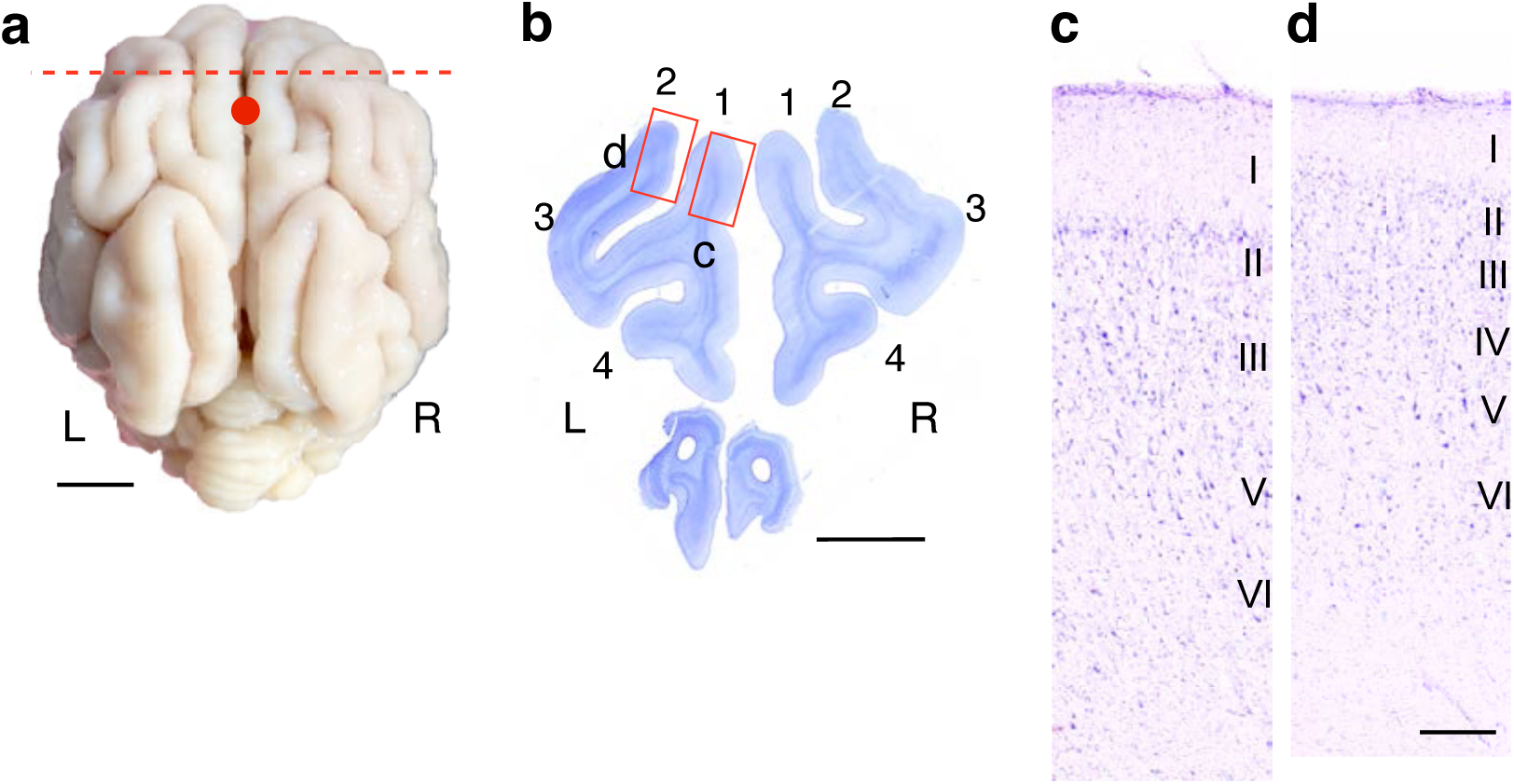
Symmetry in neocortical areas of the frontal and anterior part of the parietal lobe. (a) Dorsal view of the MMP brain with reference point, point-FS (red circle). The red dashed line (−5.04 mm anterior from point-FS) indicates the position of the coronal section corresponding to panel b. (b) Coronal brain section at −5.04 mm anterior from point-FS in panel a. Numbers represent the count of gyri in each hemisphere, indicating symmetry in the frontal and anterior part of the parietal lobe. Red rectangles indicate the regions in panel c and d, respectively (c, d). Higher magnification images of the neocortical layers (I-VI) from the corresponding regions indicated in panel b. Cortical layers are labeled on the right side of each image. Left and right sides are indicated as L and R, respectively. Scale bar for external appearance: 1 cm, Nissl staining of whole brain: 1 cm, Nissl staining with high magnification: 0.5 mm.

### Asymmetrical gyration of perirhinal lobe and cingulate cortex in frontal part of the neocortex

Next, we examined a section at +*5.58* mm from point-FS (Fig. 8a). The striatum was observed in the ventral part of the section. Wide lateral ventricules were observed medially to the striatum, whereas the insular cortex was located laterally. Interestingly, we found that gyration pattenrs are different between left and right side, and as if an ‘extra gyrus’ seems to be present in the left neocortex, dorsal to the insular (shown as ‘7’ in Fig. 8b). Combined with cingulate, frontal and parietal lobe, the left neocortex contains seven gyri, whereas six gyri were presented in the right neocortex (indicated in Fig. 8b by a number). Judging from sagittal images (Fig. 2e and Fig. 4a), the ‘extra gyrus’ on the left cerebral cortex belongs to the perirhinal cortex. The MRI images of perirhinal cortex is also shown in Suppomentary Fig. 1. The distinct pattrens of gyration was observed in sequential MRI images.

**Figure 8.**
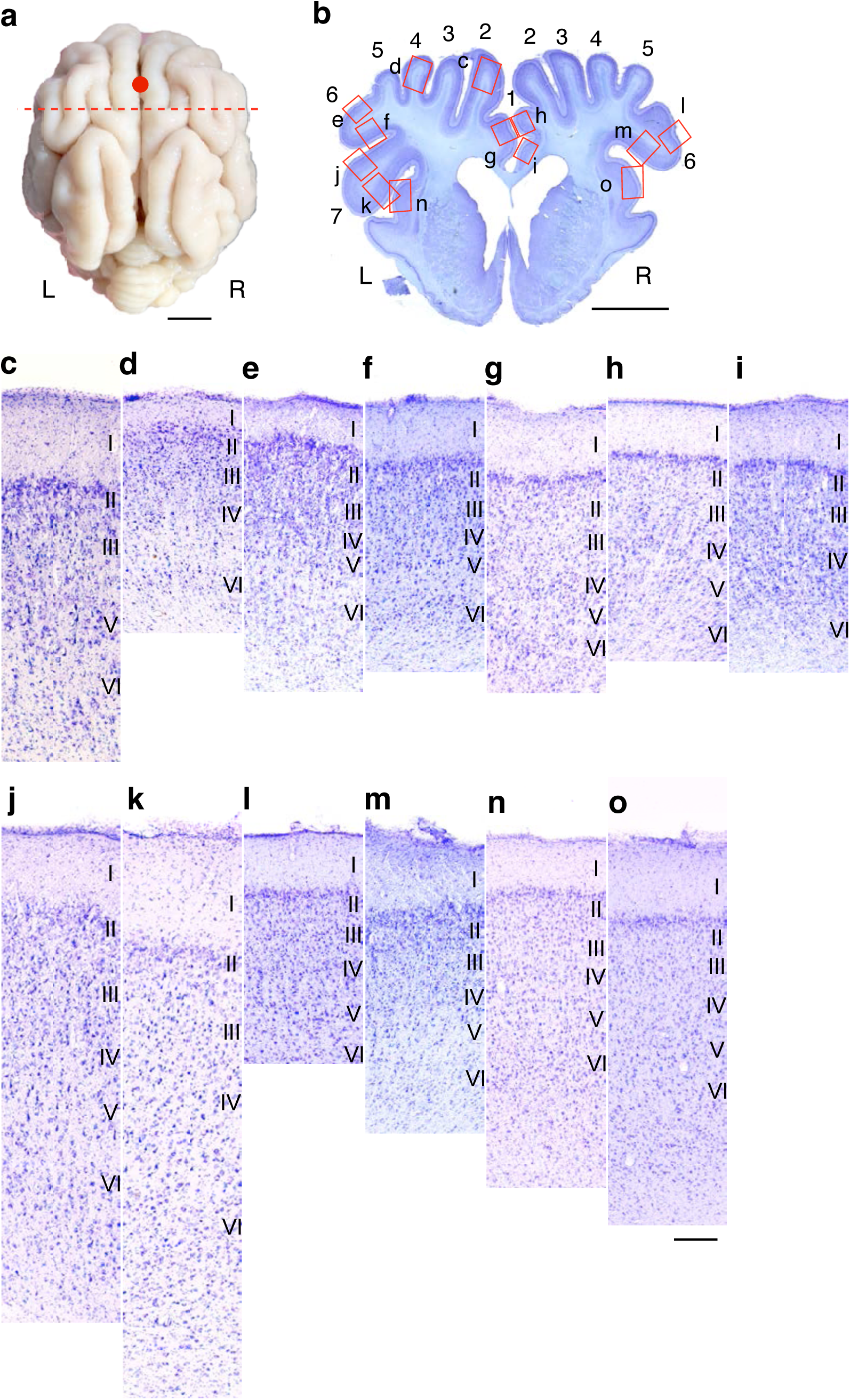
Asymmetry in neocortical areas of the anterior part of the temporal lobe. (a) Dorsal view of the MMP brain with reference point, point-FS (red circle). The red dashed line (+5.58 mm posterior from point-FS) indicates the position of the coronal section corresponding to panel b. (b) Coronal brain section at +5.58 mm posterior from point-FS in panel A. Numbers represent the count of gyri in each hemisphere, indicating asymmetry in the anterior part of the temporal lobe. Red rectangles indicate the regions in panel c to m, respectively. (c-o) Higher magnification images of the neocortical layers (I-VI) from the corresponding regions indicated in panel b. Cortical layers are labeled on the right side of each image. Left and right sides are indicated as L and R, respectively. Scale bar for external appearance: 1 cm, Nissl staining of whole brain: 1 cm, Nissl staining with high magnification: 0.5 mm.

The cytoarchitecture of the neocortical regions in the section was also examined. The middle part of the frontal lobe (Fig. 8b, indicated as ‘2-3’) clearly indicates an agranular pattern of the architecture with a large pyramidal cells in layer V, indicating the motor cortex (Fig. 8c). The middle part of the parietal lobe (Fig. 8b, indicated as ‘4’) represents strikingly different cytoarchitecture; the layer V was very thinner compared to that in frontal lobe, implicating the somatosenrory cortex (Fig. 8d). The lateral part of the parietal lobe (Fig. 8b, indicated as ‘6’) exhibited granular cytoarchitecture in both lateral and medial part (Figs. 8e, f), indicating the parietal-association area. The cingulate gyrus, which showed asymmetrical macroscopic structure as seen in Fig. 8d, exhibited a granular pattern of the cytoarchitecture (Figs. 8g-i). The ‘extra gyrus’ on left side of the cortex (Fig. 8b, indicated as ‘7’) showed distinct cytoarchitecture between lateral and medial side of the gyrus; a granular cytoarchitecture in the lateral side (Fig. 8j) and an aguranular cytoarchitecture with a relative inconspicuous layer V in the medial side (Fig. 8k), indicating that this region corresponding to the structure of perirhinal cortex. The cytoarchiteture of the right side cortex (incicated as ‘6’) exhibits similar pattens. Though neocortical layer thickness was different from that of left side, the cytoarchitecture is similar. The lateral side (Fig. 8l) have a granular cytoarchitecture, whereas medial side (Fig. 8m) shows an aguranular architecture.. The insular cortex on the left side (Fig. 8n) and on the right side (Fig. 8o) also show indistinguishable granular cytoarchitecture.

### Asymmetry of the anterior cingulate gyrus

As mentioned above, we found protrusion of right and invagination of left side of the cingulate gyrus. Next, we examined how the asymmetry extends along anterior-posterior axis (Fig. 9a). MRI images revealed that this asymmetrical structure were observed in the genu of corpus callosum (Fig. 9c) and anterior one third of the corpus callosum (Fig. 9d). In contrast, the neocortical asymmetry is hardly visible at both the frontal lobe (Fig. 9b) and the posterior part of the corpus callosum (Fig. 9e). These data indicate that neocortical asymmetry is limitedly observed in anterior, but not in posterior cingulate gyrus.

**Figure 9.**
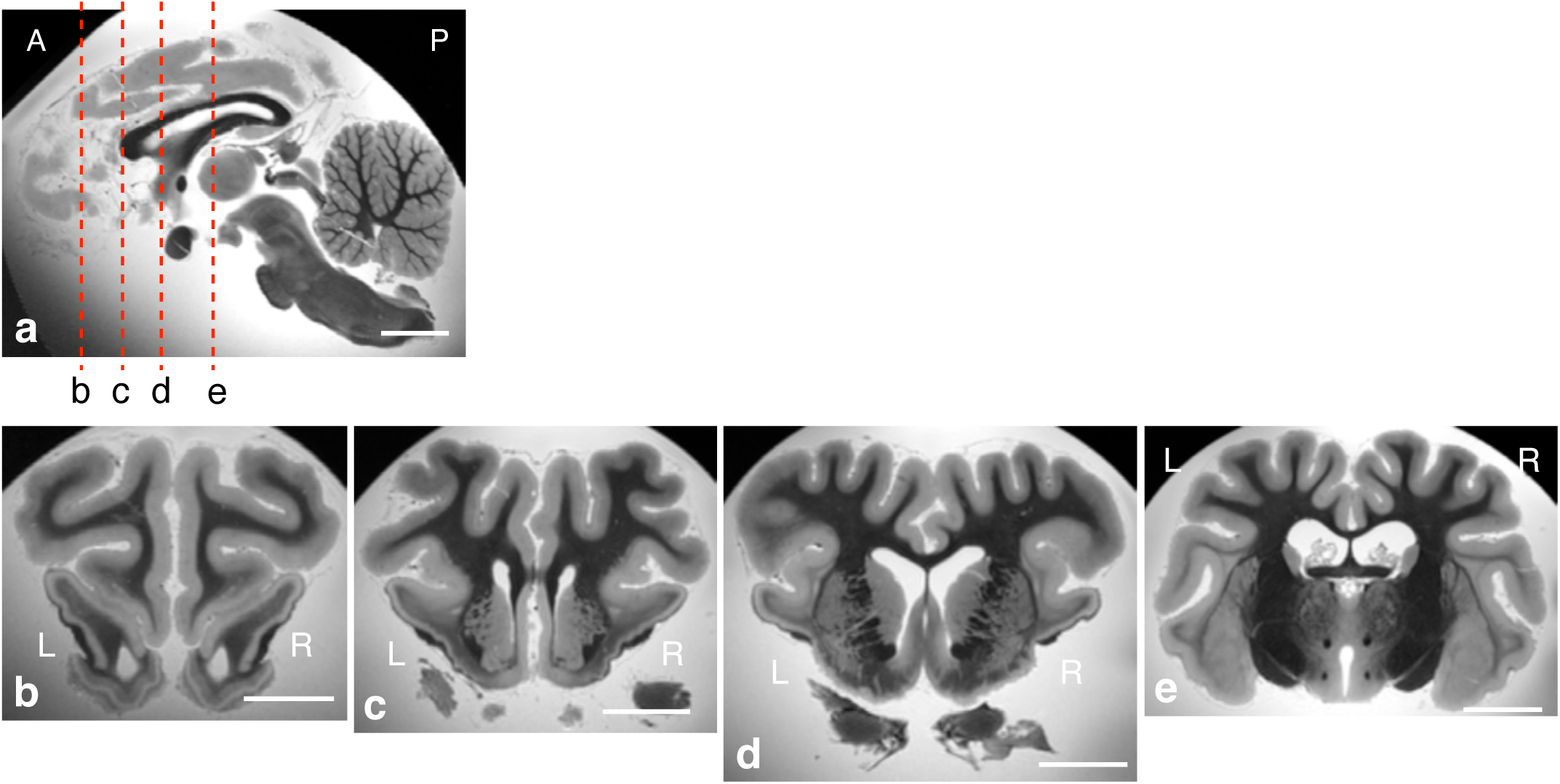
Asymmetry of the anterior cingulate gyrus. (a) Representative mid-sagittal section of T2w MRI image. Vertical dashed lines indicate the approximate plane of coronal levels used for subsequent MRI analyses along anterior-posterior axis. (b-e) Representative coronal section of T2w MRI image corresponding to vertical dashed lines. Left and right sides are indicated as L and R, respectively. Scale bar: 1 cm.

### Neocortical structure at the coronal sections of dorsal hippocampus

Given the left-right asymmetrical structure of Cer, we are interested in the neocortical structures at the level of dorsal hippocampus (+*23.42* mm from point-FS, Fig. 10a) because the asymmetrical structure is not ubiquitously found in MRI image sections. As shown in Fig. 10b, the coronal section of the MMP brain in the dorsal hippocampus also revealed another hippocampal section in the ventral part of the brain. This ventral part of the hippocampus in the section is directly connected to the entorhinal cortex. The view of the occipital cortex at this section level is symmetrical. This result indicates that neocortical asymmetry is found in special part of the neocortical areas.

**Figure 10.**
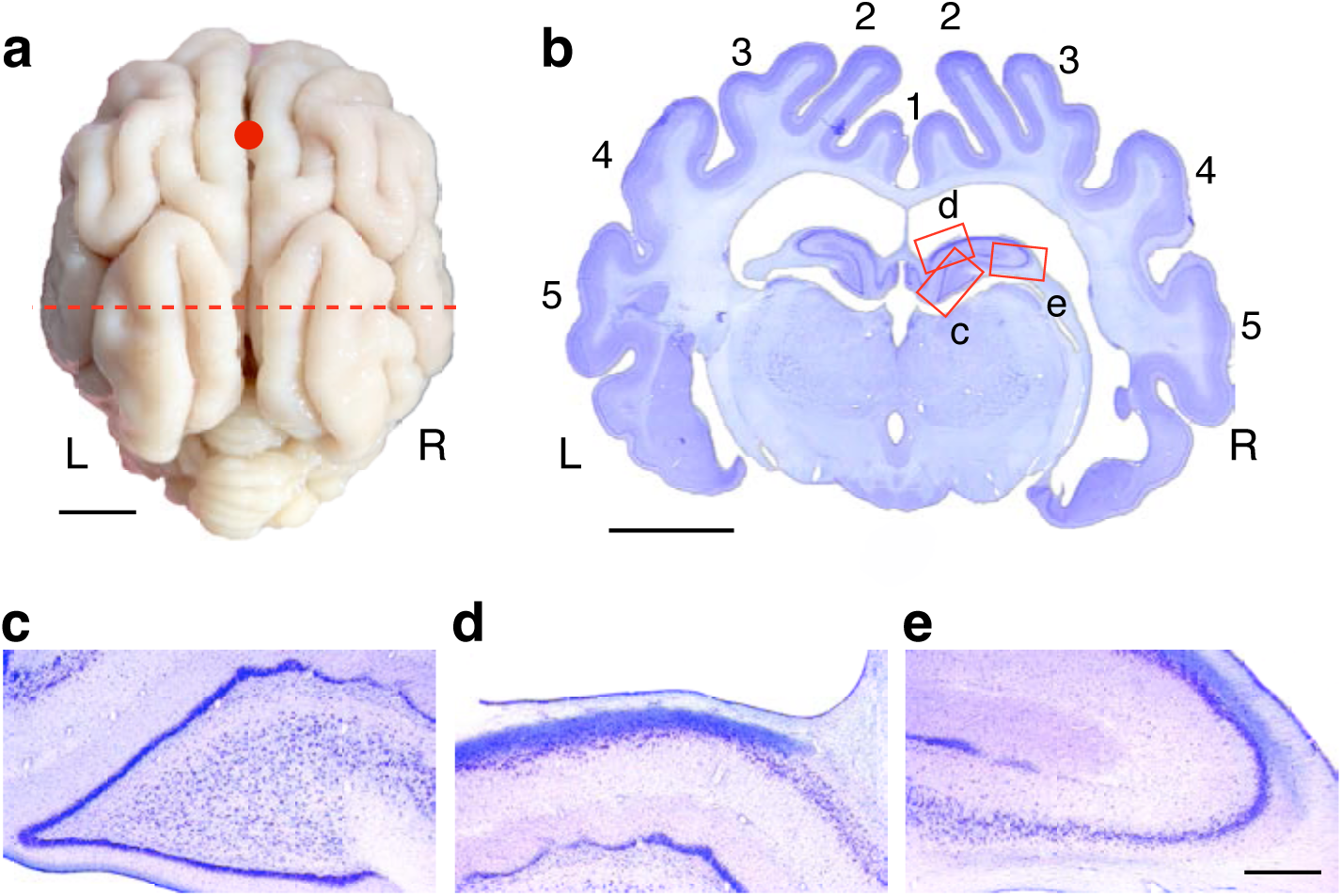
Symmetry in neocortical areas of the occipital lobe including dorsal hippocampus. (a) Dorsal view of the MMP brain with reference point, point-FS (red circle). The red dashed line (+23.42 mm posterior from point-FS) indicates the position of the coronal section corresponding to panel b. (b) Coronal brain section at +23.42 mm posterior from point-FS in panel A. Numbers represent the count of gyri in each hemisphere, indicating symmetry in the anterior part of the temporal lobe. Red rectangles indicate the regions in panel c to e, respectively (c-e). Higher magnification images of the dentate gyrus (c), CA1 (d), and CA3 (e) of the hippocampus from the corresponding regions indicated in panel b. Left and right sides are indicated as L and R, respectively. Scale bar for external appearance: 1 cm, Nissl staining of whole brain: 1 cm, Nissl staining with high magnification: 0.5 mm.

The magnified views of the dorsal hippocampus are shown in Figs. 10c, 10d and 10e. The granule cell layer of the dentate gyrus is remarkably visualized by Nissl staining and the layer forms wavy structures (Fig. 10c). The CA1 pyramidal layer indicates clear layered structure as rodents and also shows wave-like pattern (Fig. 10d). The CA3 pyramidal layer shows large pyramidal neurons which is the hallmark of CA3 area (Fig. 10e).

### Neocortical structure at the coronal sections of ventral hippocampus

Next, we observed more ventral part of the brain (+*27.00* mm from point-FS, Fig. 11a). The section contains middle-posterior part of the occipital cortex and posterior part of the hippocampus, and brain stem (Fig. 11b). The specimen contains longitudinal section of the hippocampus. The magnified views of CA3 region (Fig. 11c), CA1 region (Fig. 11d), and dentate gyrus (Fig. 11e) of the ventral hippocampus are ^39^ shown, respectively. Basic cytoarchitecture of the ventral hippocampus is similar to that of the dorsal part. However, in the dentate gyrus of the ventral part, the clear layer structure has disappeared.

**Figure 11.**
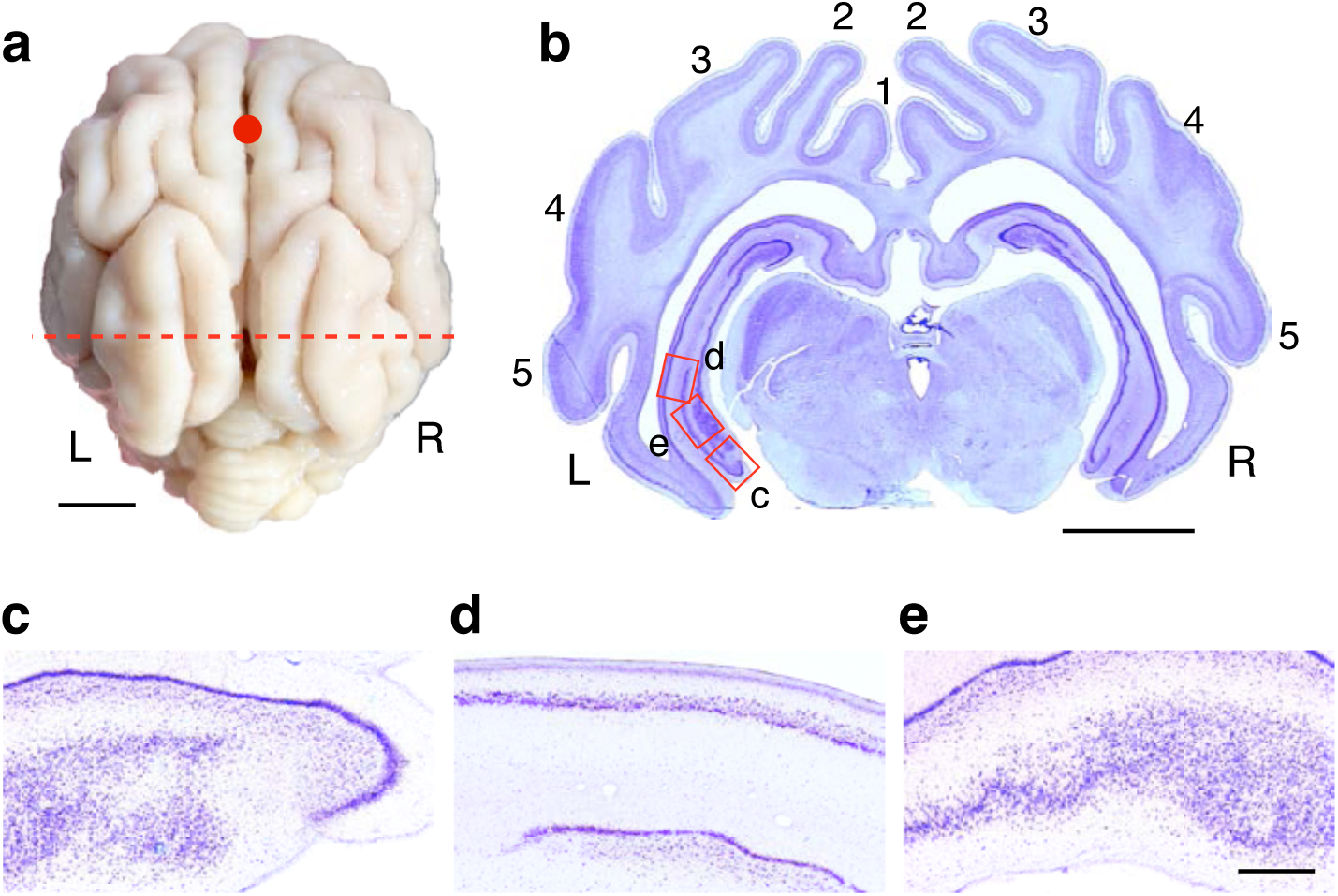
Symmetry in neocortical areas of the occipital lobe including ventral hippocampus. (a) Dorsal view of the MMP brain with reference point, point-FS (red circle). The red dashed line (+27.00 mm posterior from point-FS) indicates the position of the coronal section corresponding to panel b. (b) Coronal brain section at +23.42 mm posterior from point-FS in panel A. Numbers represent the count of gyri in each hemisphere, indicating symmetry in the anterior part of the temporal lobe. Red rectangles indicate the regions in panel c to e, respectively. (c-e) Higher magnification images of the dentate gyrus (c), CA1 (d) and CA3 (e) of the hippocampus from the corresponding regions indicated in panel b. Left and right sides are indicated as L and R, respectively. Scale bar for external appearance: 1 cm, Nissl staining of whole brain: 1 cm, Nissl staining with high magnification: 0.5 mm.

These results indicate that brain clear morphological asymmetry of MMP brain is observed only in anterior part of the neocortex and cerebellum.

## Discussion

In this report, we investigated MMP brain mesoscopic structure by slice preparation and MRI images. We prepared Nissl-stained coronal sections of the whole MMP brain and examined mesoscopic structures with the aid of MRI imaging. Previous studies using pig brain slices have assumed that pig neocortex is symmetrical along left-right axis ^38,40^, so such literature exhibited one hemisphere of the brain in sections. However, we demonstrated that cerebellum, perirhinal and anterior cingulate gyri are asymmetrical to the body axis. The asymmetry was observed similarly in both male (Nissl staining) and female (MRI image) MMPs. In contrast to slice anatomical experiments, MRI studies on pig brains have clearly classified the left and right side ^35,39^ because MRI imaging studies always describe the side of the brain rigidly. However, these reports did not focus on the asymmetrical nature of the pig brain. Moreover, compared with microscopic anatomy, MRI studies still suffer from lower spatial resolutions and need other additional exanimation methodology to definitively conclude the results. We consider that the integration of MRI imaging and slice-based microscopic analysis is essential for this research.

The asymmetry of the cerebellar vermis and paravermis has been well recognized in cats from early days ^24^. However, the physiological meaning of cat cerebellar asymmetry has not been well studied. As recent physiological experiments using cats are becoming nearly impossible, MMP would be an alternative to explore the physiological meanings of cerebellar laterality of the cerebellum. Interestingly, functional asymmetry of the cerebellum has been recently recognized in humans. The right cerebellar cortex is implicated in social recognition and language processing in humans^25,26^, and right cerebellar damage is reported to be associated with impaired language skill in human children ^25^. MMP cerebellar asymmetry is likely to be a novel animal model for studying the cerebellar function.

Among neocortical lobes, we identified apparent left-right asymmetry in the anterior cingulate gyrus. Interestingly, although asymmetrical morphology of the cingulate gyrus is not identified in mice, hippocampal asymmetry has been well documented ^10,11,13^, and the cingulate gyrus and hippocampus have reciprocal projections ^27,41^. Out finding hints the exsitance of hippocampal asymmetry in MMPs. Hippocampal left-right asymmetrical function is also recognized in humans ^42^. In contrast, we did not detect apparent asymmetry in entorhinal cortex, which is the principal target of hippocampal synaptic output and major input. To investigate possible left-right differences of the hippocampal functions in MMPs, future diffusion tensor imaging studies and trasce studies are expected to clarify left-right differences in neural circuitry between anterior cingulate cortex and hippocampus.

We also identified distinct pattens of gyration between left and right perirhinal cortex. The location of asymmetry is just next to the border of perirhinal and parietal cortex, and the most ventral part of the neocortical areas. Insular cortex is on the opposite and inner side of this gyrus. The cytoarchitecture of this gyrus exhibited large pyramidal cells in layer V in lateral side, but not in medial side. The data suggests the gyri in ventral extreme of the perhichinal cortex (incidated as ‘7’ on the left side and ‘6’ on the right side) project to lower part of the brain. This asymmetry suddenly disappears in the more posterior part of the neocortex (the border is around the middle part of the corpus callosum). Since perirhinal cortex feeds neocortical information to hippocampus directly and via entorhinal cortex^28,30^, the area is also implicated in hippocampal memory formation ^31,43^ In summary, we report that the MMP brain has clear mesoscopic left-right asymmetry in several areas of the brain. The identified asymmetrical areas hint that the MMP brain asymmetry would be functionally correlated to brain asymmetry that has been recognized in rodents and primates. We propose that MMP brains may be one of the valuable animal models and help us understand brain asymmetry in future studies.

## Author contributions

YS conceptualized the experiments. YF, YS, KY, RM, TA, NW, AS, MT, and NG performed the experiments. KY designed the figures. YS and KY wrote the manuscript. SC and AN supervised the experiments.

## Acknowledgement

We are gratefule to President Noriaki Satake (Fuji Micra Inc.) for kind donation of a microminipig.

---

**Supplementary Figure 1.**
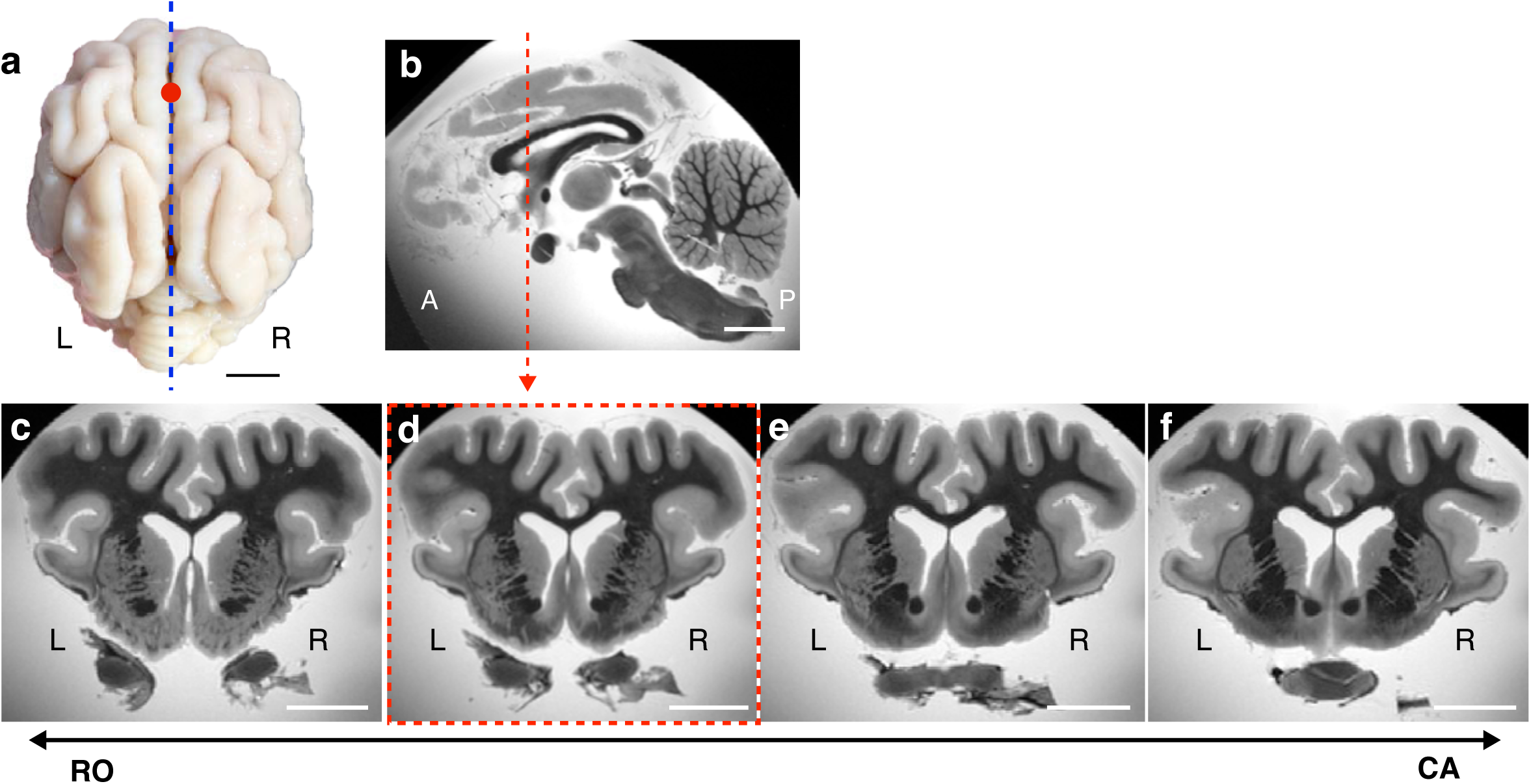
Asymmetrical gyration of the perirhinal cortex observed in MRI images. (a) Dorsal view of the MMP brain with reference point, point-FS (red circle). The blue dashed line indicates the midline. (b) Representative mid-sagittal section of T2w MRI image. Vertical dashed lines indicate the approximate plane of coronal levels cooresponding to panel d. (c-f) Representative coronal section of a T2w MRI images arranged from anterior to posterior. Left and right sides are indicated as L and R, respectively. Scale bar: 1 cm.

